# All-optical interrogation of brain-wide activity in freely swimming larval zebrafish

**DOI:** 10.1101/2023.05.24.542114

**Authors:** Yuming Chai, Kexin Qi, Yubin Wu, Daguang Li, Guodong Tan, Yuqi Guo, Jun Chu, Yu Mu, Chen Shen, Quan Wen

## Abstract

We introduce an all-optical technique that enables volumetric imaging of brain-wide calcium activity and targeted optogenetic stimulation of specific brain regions in freely swimming larval zebrafish. The system consists of three main components: a 3D tracking module, a dual color fluorescence imaging module, and a real-time activity manipulation module. Our approach uses a sensitive genetically encoded calcium indicator in combination with a long Stokes shift red fluorescence protein as a reference channel, allowing the extraction of Ca^2+^ activity from signals contaminated by motion artifacts. The method also incorporates rapid 3D image reconstruction and registration, facilitating *real-time* selective optogenetic stimulation of different regions of the brain. By demonstrating that selective light activation of the midbrain regions in larval zebrafish could reliably trigger biased turning behavior and changes of brain-wide neural activity, we present a valuable tool for investigating the causal relationship between distributed neural circuit dynamics and naturalistic behavior.

**Highlights:**

- We develop an all-optical technique that enables simultaneous whole brain imaging and optogenetic manipulation of selective brain regions in freely behaving larval zebrafish.
- A combination of a genetically encoded calcium indicator and a long Stokes-shift red fluorescence protein, together with the adaptive filter algorithm, enables us to reliably distinguish calcium activity from motion-induced signal contamination.
- Rapid 3D image reconstruction and registration enables real-time targeted optogenetic stimulation of distinct brain regions in a freely swimming larval zebrafish.

## 1 Introduction

One of the central questions in systems neuroscience is understanding how distributed neural activity over space and time gives rise to animal behaviors (1, 2). This relationship is confounded by recent recordings in several model organisms, which reveal that brain-wide activity is pervaded by behavior-related signals (3–7). All-optical interrogation, which enables simultaneous optical read-out and manipulation of activity in brain circuits, opens a new avenue to investigate neural dynamics that are causally related to behaviors and neural representation of behaviors that are involved in different cognitive processes (8–11).

All-optical neurophysiology has been successfully applied to probe the functional connectivity of neural circuit *in vivo* and the impact of genetically or functionally defined group of neurons on the behaviors of head-fixed animals (12–14). Here we extend this technique to *freely swimming* larval zebrafish, which allows simultaneous targeted stimulation of the brain region of interest and read-out of whole brain Ca^2+^ activity during naturalistic behavior, such that all sensorimotor loops remain intact and active. Our approach leverages recent advancements in volumetric imaging and machine learning: (1) with the advent of light-field microscope (LFM), brain-wide neural activity can be captured rapidly and simultaneously; (2) deep neural network-based image detection and registration algorithms enable robust real-time tracking and brain region selection for activity manipulation.

There have been several reports on whole brain imaging of freely swimming zebrafish (15–20). However, a serious problem can hinder the wide use of this technique: swimming itself causes substantial fluctuations in the brightness of the neural activity indicator. These fluctuations can interfere with the accurate interpretation of true neural activity in zebrafish (19). In an effort to overcome this issue, we have integrated an imaging channel designed for simultaneous panneuronal imaging of a long Stokes shift and activity-independent red fluorescence protein (Fig. S1). This protein shares the same excitation laser as the Ca^2+^ indicator. The incorporation of a reference channel, alongside the implementation of an adaptive filter algorithm, enables us to correct activity signals tainted by motion artifacts resulting from the zebrafish’s swift movements.

Fig. 1 shows a schematic of our system that integrated tracking, dual color volumetric fluorescence imaging and optogenetic manipulation. We performed *simultaneous* brain-wide Ca^2+^ signal and reference signal recording using the fast eXtended Light Field Microscope (XLFM) (16). An optogenetic module was incorporated into the imaging system to enable real-time activity manipulation in defined brain regions in a freely swimming larval zebrafish.

**Figure 1.**
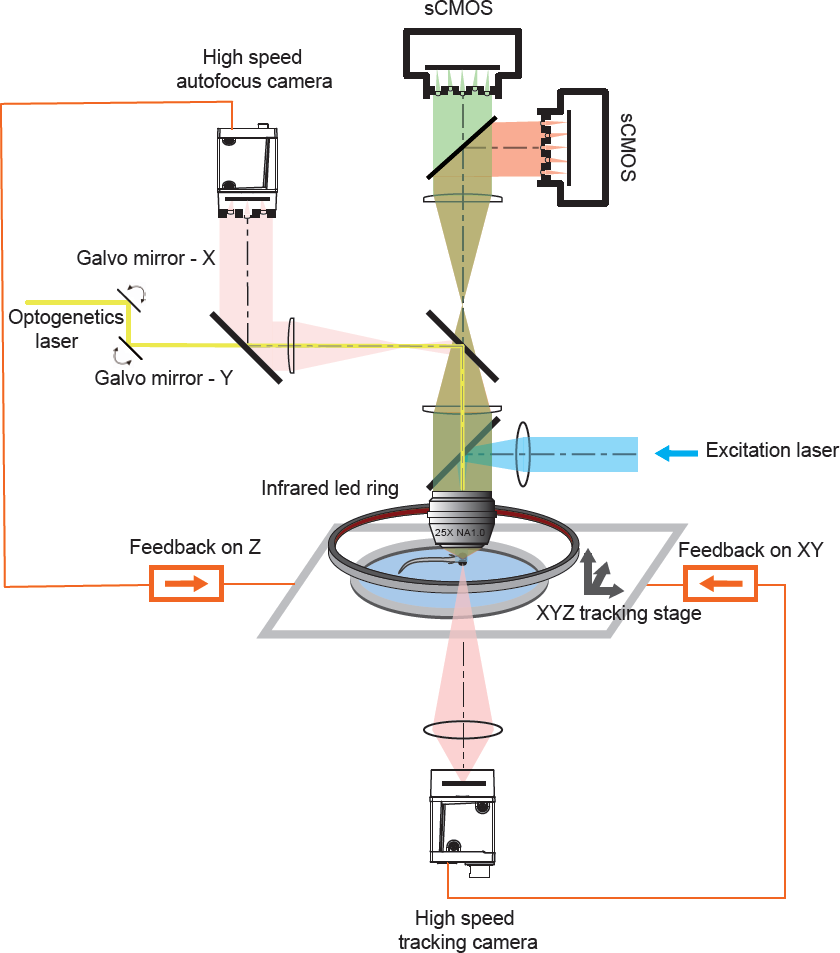
Schematics of the dual color whole brain imaging and optogenetic system. The system integrated tracking, dual color light-field imaging, and optogenetic stimulation. A convolutional neural network (CNN) was used to detect the positions of fish head from dark-field images captured by the near-infrared (NIR) tracking camera. A tracking model converted the real-time positional information into analog signals to drive the high-speed motorized stage and compensate for fish movement. The neural activity-dependent green fluorescence signal and the activity-independent red fluorescence signal were split into two beams by a dichroic mirror before entering the two sCMOS cameras separately. The dichroic mirror was placed just before the micro-lens array. Both the red and green fluorophore can be excited by a blue laser (488 nm). An x-y galvo system deflected a yellow laser (588 nm) to a user-defined ROI in the fish brain for real-time optogenetic manipulation with the aid of fast whole-brain image reconstruction and registration algorithm.

## 2 Results

### 2.1 Tracking system

To reliably maintain the head of a swimming fish within the field of view (FoV) of the microscope, we redesigned our high-speed tracking system (16) with two major changes. First, to correctly identify the position of the fish head and yolk from a complex background, we replaced the conventional computer vision algorithms for object detection (that is, background modeling and adaptive threshold) with a U-Net (21) image processing module (Methods). This approach greatly improves the accuracy and robustness of the tracking in a complex environment while ensuring a high image detection speed (< 3 ms). Second, we combined the current position of the fish and its historical motion trajectory (17) to predict fish’s motion (Fig. 2a). This allows the system to preemptively adjust the stage position to keep the fish in view, even when it is swimming quickly.

**Figure 2.**
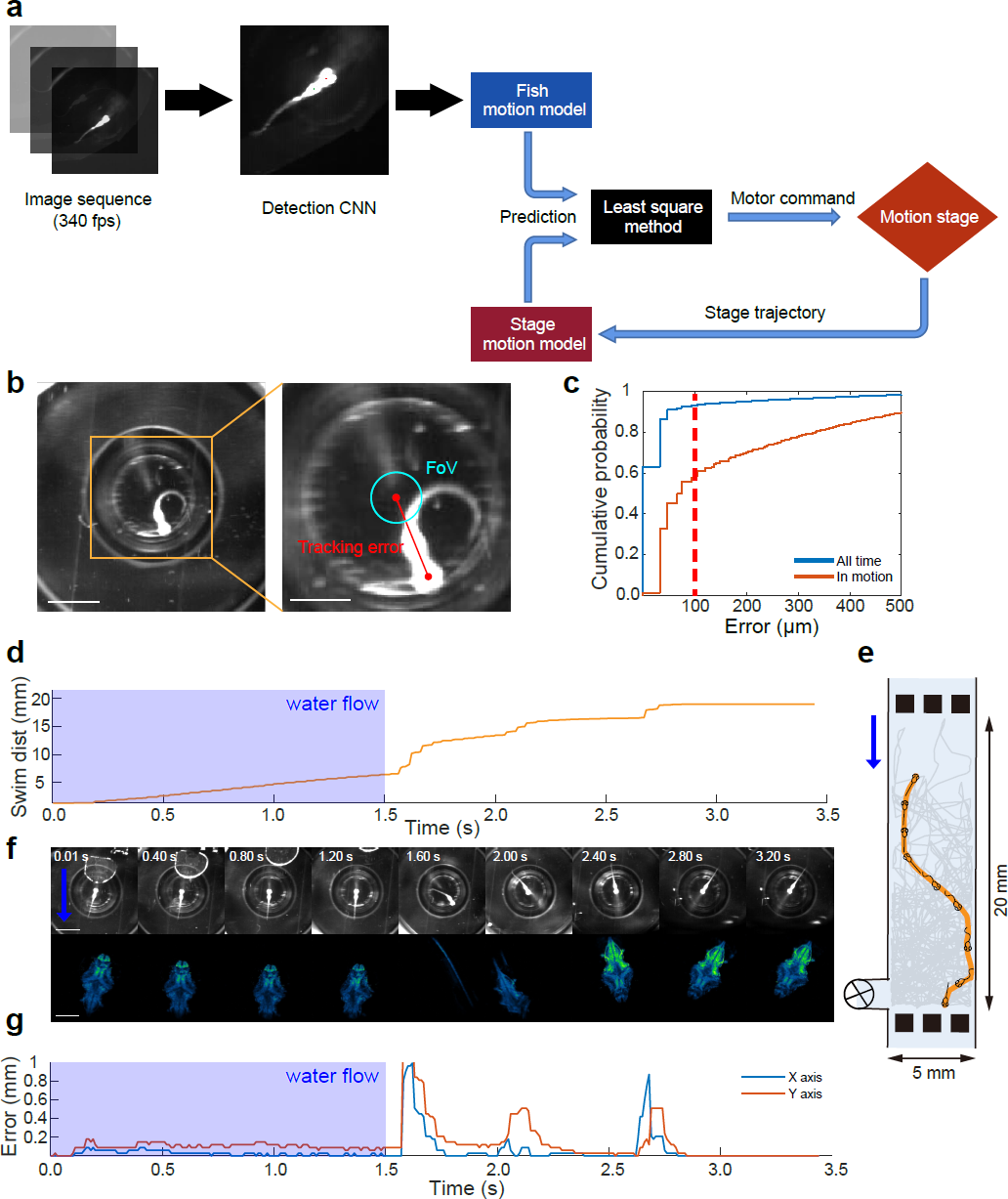
Tracking system. **a.** Flowchart of the tracking system. **b.** A near-infrared (NIR) image captured by the tracking camera, with the tracking error highlighted. The cyan circle indicates the field of view (FoV) of the XLFM. The tracking error is the distance between the center of the fish head and the center of the FoV. The scale bar is 2.5 mm (left) and 1 mm (right). **c.** The cumulative distribution of the tracking error, based on all time points or when the fish was in motion. "In motion" is defined as a tracking error that lasts for at least 30 ms and exceeds 50 *µ*m in maximum magnitude. The data comes from 34 fish, each of which was tracked for more than 10 minutes. The red dashed line is the maximum tolerable error, the distance beyond which the fish brain is not completely visible in the FoV. **d-g** Tracking example of a freely swimming larval zebrafish stimulated by water flow. **d.** Swimming distance during example trajectory. **e.** The example trajectory. The yellow line indicates the example trajectory of the fish, and the gray lines indicate all other movements in the microfluidic chip. **f.** Top, NIR tracking video images. The scale bar is 2.5 mm. Bottom, reconstructed whole brain fluorescence images obtained during this trajectory. Scale bar, 300 *µ*m. **g.** Tracking error during this example trajectory. Note that the applied water flow (arrow in **e**) forced the fish to move backward (VideoS1), an unexpected movement pattern for the motion prediction model. As a result, the system shows a larger tracking error in the blue-shaded period.

To quantify the tracking performance, we define the tracking error as the distance between the center of the fish head and the center of the microscope FoV (Fig. 2b right). The FoV of the XLFM is 800 µm in diameter, and the size of brain along the rostrocaudal axis is about 600 µm. Therefore, a tracking error less than 100 µm is sufficient to capture the image of the entire brain. We tested our tracking system in several experimental paradigms, including spontaneous behavior, swimming in the presence of water flow, and during optogenetic stimulation. We found that about 92.6% of the frames were within 100 µm tracking error at all times (a total of 13,801,838 frames, Fig. 2c). During locomotion, 58.69% of the frames were within this range of tracking error (n = 26,930 bouts, 2,293,258 frames).

Fig. 2d-g shows the performance of the tracking system when a water flow stimulus was applied to the fish. On this occasion, a large bubble appeared in the microfluidic chamber (Fig. 2f, top). Despite a distracting background, the image detection module was still able to accurately identify the position of the fish and keep the head of the fish within the microscope FoV (Fig. 2f, bottom). Taken together, these results demonstrate that our tracking system is highly reliable and can be used in a variety of behavior experiments.

### 2.2 Dual color volumetric image alignment

Two-color fluorescence imaging (Fig. 1) enables us to use a reference signal (Fig. S1) to correct for Ca^2+^ signal artifacts caused by zebrafish movements. Fig. 3a shows each step in the image processing pipeline. Briefly, the reconstructed 3D volumetric image frames of the activity-independent red channel were registered and aligned with a template. The pairwise transformation matrix was then applied to the 3D images of the green channel. The

**Figure 3.**
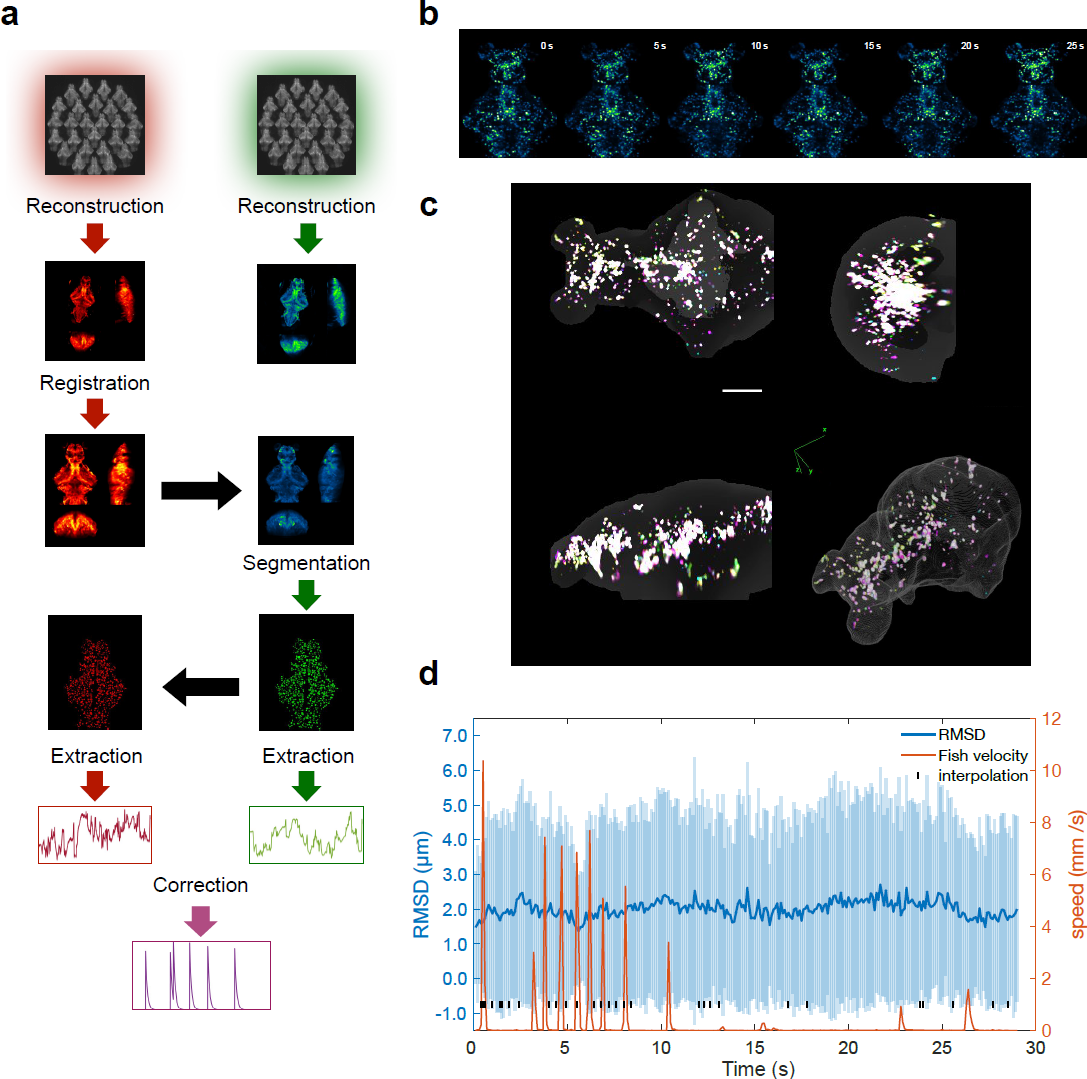
Dual-color image registration. **a.** Dual-color image processing pipeline (Methods) . The flow chart highlights the following steps: 3D light-field reconstruction; multi-scale image alignment; region of interest (ROI) segmentation; and Ca^2+^ signal extraction and correction. The black arrows indicate that identical operations can be directly applied to a different channel. **b.** Aligned images of sparsely labelled EGFP zebrafish, guided by the reference channel (VideoS2). **c.** 3D visualization of the alignment of **b**. **d.** The root-mean-square displacement (RMSD) of neuronal center of mass positions across time. This displacement was measured in relation to their coordinates within a specified reference frame and was averaged across all neurons that could be matched within a frame; the shaded region indicates SD. The red curve shows the instantaneous swimming speed of larval zebrafish.

aligned green channel images were segmented into region of interest (ROI) based on the spatiotemporal correlation of the intensity of the voxel (Methods), and the Ca^2+^ signal in each ROI was extracted and corrected.

Aligning whole brain image frames is one of the critical steps in extracting the Ca^2+^ signal accurately. However, the brain regions could deform significantly and the fluorescence intensity of most ROIs could change significantly in swimming zebrafish. These factors make whole-brain image registration a challenging task.

Here we used CMTK toolkit (22) and imregdemons function in MATLAB (23) to complete the rigid and non-rigid registration of a 3D image, respectively (Methods). To test our alignment algorithm, we injected the Tol2-elavl3:h2b-EGFP plasmid into fertilized eggs of elavl3:h2b-LSSmCrimson zebrafish. Due to the uneven distribution of the plasmid in the eggs, EGFP did not achieve whole brain neuronal expression in this generation of zebrafish, but rather showed expression patterns with different degrees of sparseness. We selected zebrafish with moderately sparse expression of EGFP at 6 dpf and recorded their dual-channel images during fish movement. Due to the sparse expression of EGFP, individual neurons randomly distributed in different brain regions can be seen in the 3D reconstructed images. After registering red channel frames with pan-neuronal expression of LSSmCrimson (Fig. 3a), we applied the same transformation to the sparse EGFP images (Fig. 3b). This allowed us to view the alignment results for each neuron in the green channel (Methods) (VideoS2), and to test and optimize our registration algorithm.

Fig. 3c represents the aligned EGFP multi-frame images (Fig. 3b) in different colors, overlaid to visualize the alignment effect. Whiter colors indicate better alignment (Fig. 3c). The sparsity of EGFP expression allows us to track about 160 neurons over time according to the spatiotemporal co<u>ntinuity o</u>f an object (Methods and Fig. S2). We quantified the root-mean-square displacement (RMSD), namely 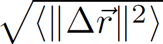, of the center of mass of a neuron relative to its coordinates in the reference frame (Fig. 3d), where the blue line indicates an average over all the neurons whose correspondents could be identified. The RMSD (Fig. 3d) is much smaller than the ROI size (8.8-18.3 µm, 25% - 75% quantile) resulting from our segmentation algorithm (Methods). Together, these results suggest that our data processing pipeline is effective in aligning whole brain images of larval zebrafish and thus can extract Ca^2+^ signals from most ROIs accurately.

### 2.3 Motion artifact correction

Rapid movements of larval zebrafish (translation and tilt) within FoV can cause significant changes in fluorescence intensity in both green and red channels, even when neural activity does not change (19). Here, we introduce an adaptive filter (AF) algorithm (Fig. 4a) to correct for motion artifacts (Methods). We use the AF algorithm to dynamically predict the green signal *ĝ*(*n*) from the signal in the red channel so that the difference between the predicted and the actual green signal in the current time frame n, e(n) = *ĝ*(*n*) − g(n), is as small as possible. In the absence of neural activity, e(n) is expected to fluctuate around 0. When there is neural activity, the resulting Ca^2+^ signal would rise rapidly and e(n) would have a large positive value. We identified the predicted *ĝ*(*n*) as the time-dependent baseline of the signal in the green channel due to zebrafish movements. The Ca^2+^ activity was inferred from the normalized signal difference ^(^g(n) − *ĝ*(*n*)^/^/*ĝ*(*n*).

**Figure 4.**
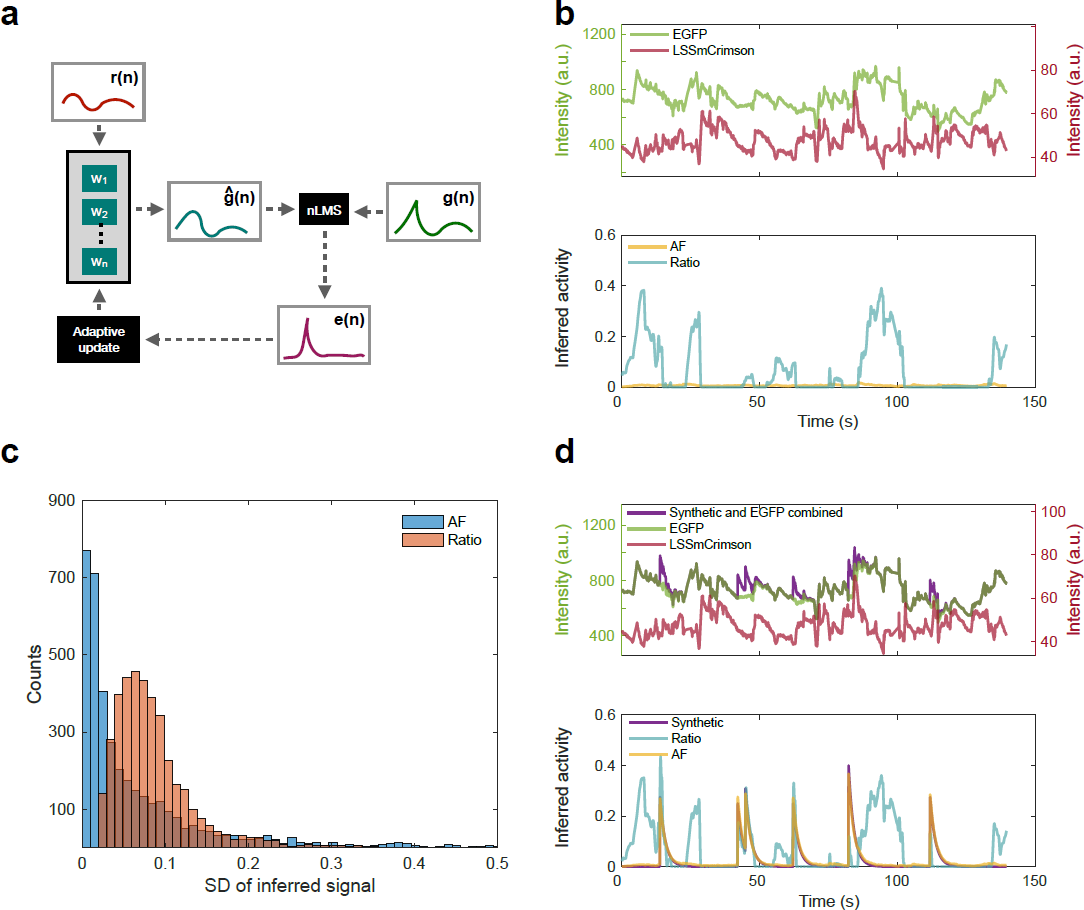
AF algorithm for signal correction. **a.** Schematics of the AF algorithm. The AF algorithm works by first estimating the motion artifacts in the green channel *ĝ*(*n*) using a weighted sum of the red channel signal r(n) over a recent history. The weights w are dynamically updated so that the residual error, e(n) = g(n) - *ĝ*(*n*), is minimized. **b.** Top, representative raw fluorescence signals from an ROI in freely swimming zebrafish with panneuronal expression of EGFP and LSSmCrimson. Bottom, inferred activity traces. The AF method is more accurate than the conventional ratiometric method (Ratio) in removing motion artifacts. **c.** The histogram shows the distribution of the inferred ROI activity level, defined as the standard deviation of activity over time. Related to **b**. **d.** Top, a raw EGFP fluorescence signal with randomly added synthetic neural activity (purple). Bottom, inferred activity trace (yellow).

The AF algorithm uses the history-dependent correlation between the green fluorescence signals (jGCaMP8s (24)) and the red fluorescence signals (LSSmCrimson) to correct for changes in motion-induced signals. These changes are caused by two major factors.

- Inhomogeneous light field. When a larval zebrafish moves, the intensity of the excitation light varies across the FoV. In this case, the change of fluorescence intensity in the green channel and that in the red channel differ by a *proportionality constant*, namely *δg*(*n*) = *α* · *δr*(*n*). A simple ratiometric division (25, 26) between the green and red channels can largely correct for this effect.
- Scattering and attenuation. When the larval zebrafish tilts its body, the fluorescence emitted from the same neuron is scattered and obscured by brain tissues and body pigments. This complicated time-varying process is likely to have a chromatic difference, leading to *disproportionate changes* in the intensity between the green and red signals. The latter effect cannot be easily corrected for by a direct division.

The history dependence of the signal comes from the observation that motion-induced fluorescence changes can persist over multiple frames.

To verify the effectiveness of the motion correction algorithm, we constructed a transgenic zebrafish line with pan-neuronal nucleus expression of EGFP and LSSmCrimson. We then simultaneously recorded the green and red fluorescence signals in a freely swimming larval zebrafish. LSSmCrimson was used to remove the fluctuation of the EGFP signal caused by animal movements. The *ideal* corrected EGFP signal, which does not change due to neural activity, would be close to 0. Fig. 4b shows representative fluorescence signals from an ROI before and after correction. The AF algorithm significantly reduced the motion-induced green fluorescence change compared to the ratiometric method. The improvement over the conventional method was quantified by plotting the distribution of the corrected signal fluctuation (i.e., the standard deviation) across all ROIs (Fig. 4c).

After demonstrating that our dual-channel motion correction algorithm can largely eliminate signal changes due to animal movements, we next show that the same algorithm can extract true Ca^2+^ signal due to neural activity. First, we overlay computer-generated randomly timed neural activity on EGFP signals (Fig. 4d, top), namelyg*^′^*(*n*) = *g*(*n*)(1 + *a*(*n*)), where *a*(*n*) is synthetic neural activity. We then asked whether the AF algorithm could correctly identify these synthetic signals (Fig. 4d, bottom) by examining the correlation coefficient *r* between a(n) and the inferred signal. We identified three factors that contribute to the precision of the AF algorithm (Fig. S5): a higher amplitude of Ca^2+^ activity, a higher correlation between the green and red channel signals, and a lower coefficient of variation (CV) of the red channel signal. We established a criteria (see Methods) for screening ROIs using the two-channel signal correlation and the CV of the LSSmCrimson signal, such that the inferred signal and the synthetic signal would exhibit a high correlation. Inferred results from brain regions that met this criterion were considered reliable and used for further analysis.

Second, we wanted to investigate whether we could identify brain regions with similar stimulus-triggered Ca^2+^ activity patterns in freely swimming larval zebrafish and in the same animal that was immobilized in agarose. We decided to use a *spatially invariant* external stimulus: the blue excitation laser (488 nm, Fig. 1). In other words, we suddenly turned on the blue excitation light when the zebrafish was in the dark, thus inducing brain activity. The main advantage of this approach is that the blue excitation light bathes on the zebrafish head in a cylindrical shape, so the blue laser remains a relatively invariant stimulus for the animal even if its body orientation changes. Furthermore, the sudden onset of blue light in the dark is a powerful stimulus that can easily trigger brain activity.

Fig. 5a shows the procedure of our blue light stimulation experiment. First, we presented a freely-swimming larval zebrafish with a 20-second blue-light stimulation (1 ms pulsed illumination at 25 Hz) followed by a 30-second dark period. The same animal was then immobilized with agarose and the same visual stimuli were applied. Brain-wide Ca^2+^ activity was recorded in both trials. We identified ROIs in immobilized zebrafish that showed prominent activity in response to the onset of repeated blue light stimulation (Fig. 5b, right). We then examined neural responses in the same ROIs when the animal was swimming freely. The similarity of the Ca^2+^ dynamics between different experimental conditions (Fig. 5a) was quantified by correlation analysis (Fig. 5d, left), where each data point represents a single trial from a single ROI. Many trials exhibited high correlation, and similarity improved after we applied the AF algorithm to remove motion artifacts.

**Figure 5.**
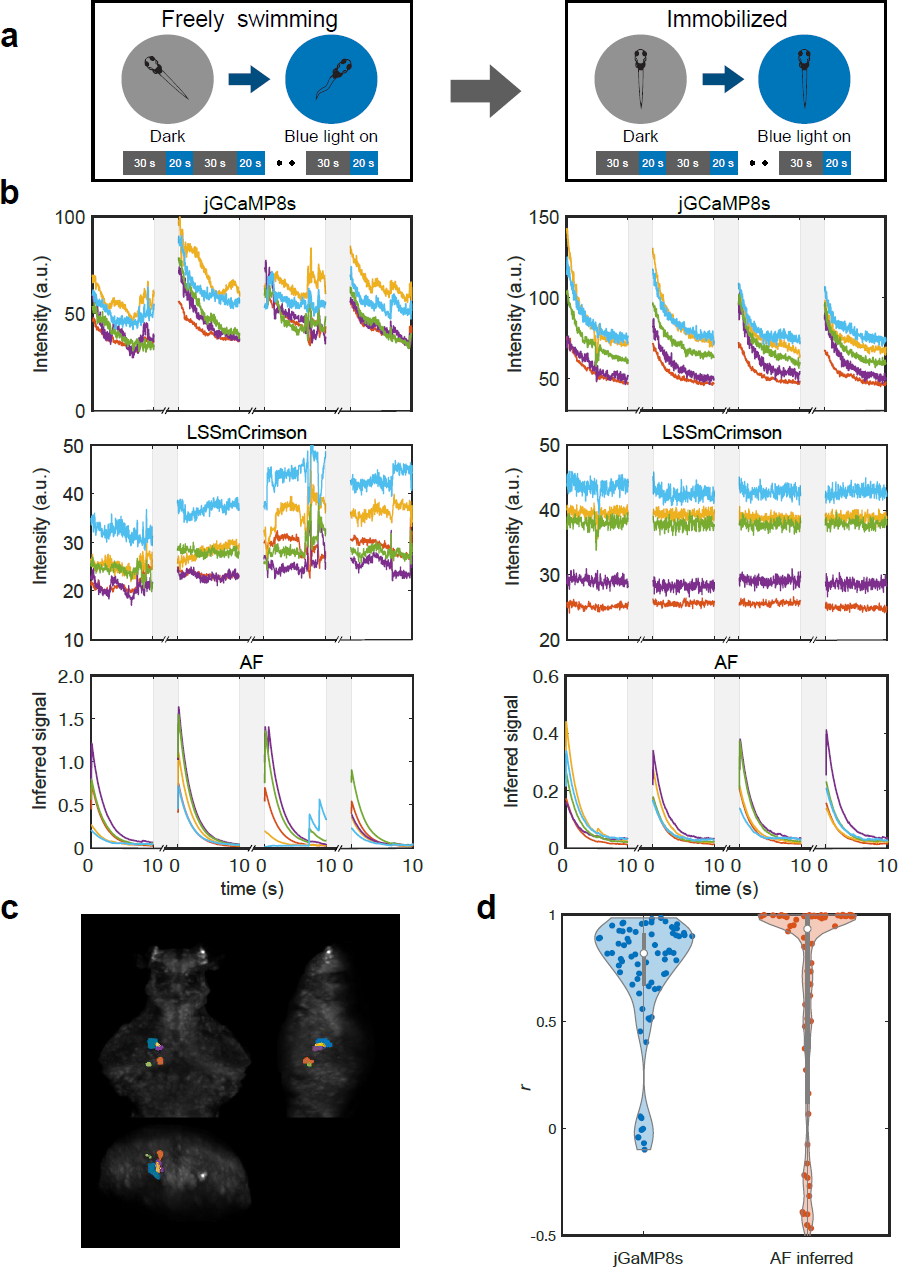
Blue light stimulation. **a.** Experimental paradigm of the blue light stimulation. A larval zebrafish was freely swimming under our tracking microscope while blue excitation light (1 ms pulsed stimulation, 2.5% duty cycle) was suddenly turned on. The light was turned on and off at 20/30 second intervals. The same animal was then immobilized in low melting point agarose and an identical pattern of light stimulation was applied. Brain-wide calcium activity was recorded in both conditions. **b.** Top: jGCaMP8s and LSSmCrimson raw fluorescence signals in 5 representative ROIs in which neurons showed prominent Ca^2+^ activity after light stimulus onset. Shaded regions indicate the dark period. Bottom: inferred calcium activity using the AF algorithm. Left panels are recordings from freely swimming condition while right panels are from immobilized condition. **c.** The spatial location of the brain regions in **b**. **d.** Violin charts of trial-to-trial pairwise correlation between Ca^2+^ activity in freely-swimming and immobile conditions. Left, correlations of raw jGCaMP8s signals; right, correlations of AF inferred signals. Each violin chart represents the distribution of *r* for each of the 4 trials (see **b**) in 18 selected ROIs (a total of 72 data points).

### 2.4 Real-time optogenetic manipulation of freely swimming larval zebrafish

After introducing the dual-color AF method to extract brain-wide Ca^2+^ activity, we now describe the optogenetic system that enables real-time manipulation of user-defined brain regions in freely behaving larval zebrafish (Fig. 1). A user first selects the region to be stimulated on the zebrafish brain browser (ZBB) atlas (Fig. 6b, left) (27). The system then translates the region into actual locations on the fish brain and delivers photo-stimulation through real-time image processing and coordinate transformation.

**Figure 6.**
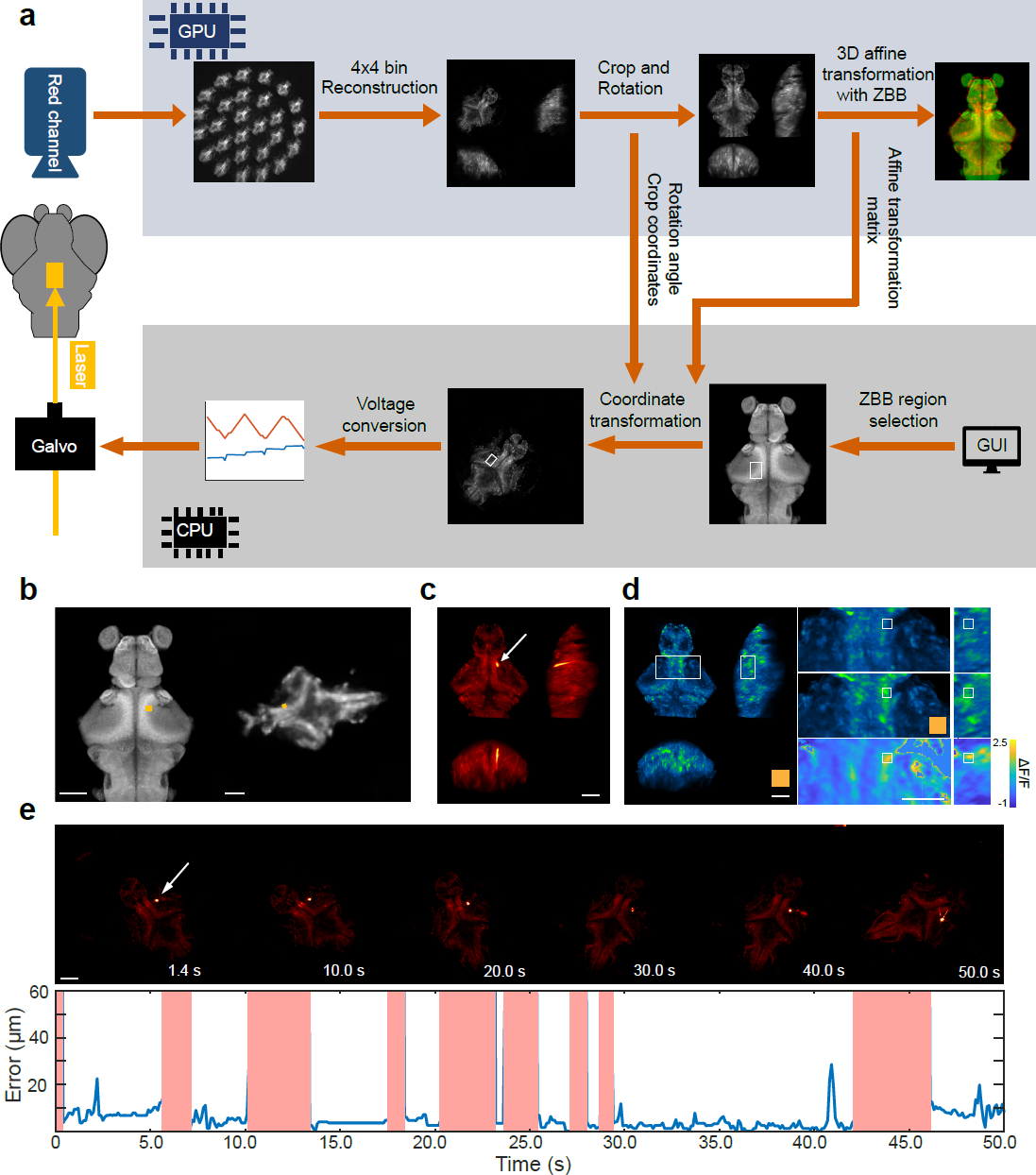
Real-time optogenetic system. **a.** Schematics of the optogenetic stimulation workflow. **b.** Transformation of a user-defined brain region on a ZBB atlas (left) to a real-time position on a freely swimming zebrafish (right). The yellow rectangle shows the stimulation pattern. Scale bar, 100 *µ*m. **c.** Pseudo-color fish brain pixel intensity averaged across 200 registered red channel fluorescence images. The white arrow indicates the location of the red fluorescence excited by the laser beam (Methods). Scale bar, 100 *µ*m. **d.** Left, green channel image. Scale bar, 100 *µ*m. Right, a zoomed-in image around the stimulated region (white rectangle) before (top) and during (middle) yellow light stimulation. The color map (bottom) indicates the change in fluorescence intensity (Methods). Scale bar, 50 *µ*m. **e.** Top, images of the fish brain during optogenetic stimulation. Scale bar: 100 *µ*m. Bottom, the displacement between the actual position of the light beam and the targeted position. We deflected the laser out of the FoV during periods of rapid fish movements, indicated by the shaded red regions.

Fig. 6a shows the workflow of the optogenetic module. We used the red channel fluorescence image for brain region selection. To achieve online image processing, we speed up the reconstruction and registration algorithm by resizing the images, reducing the number of iterations in the reconstruction algorithm (16), and using a deep neural network model to compute the affine transformation matrix (28). These optimizations reduce the image processing time to 80 ms, faster than the acquisition speed of fluorescence imaging (10 Hz). After the coordinates of the user-selected region on the ZBB atlas are translated into the coordinates on the real-time image, the position is converted into a two-dimensional analog voltage signal. This signal is used to control the rapid deflection of the Galvo mirror in the X and Y directions, which completes the optical stimulation of a specified region in the larval zebrafish brain.

Here we used transgenic zebrafish with pan-neuronal expression of jGCaMP8s (24), LSSmCrimson and the light-sensitive protein ChrimsonR (29) (elavl3:H2B-jGCaMP8s - elavl3:H2B-LSSmCrimson × elavl3:ChrimsonR-tdTomato, 7 dpf) to test the capability of our system. A minimal area was selected in the ZBB atlas (Fig. 6b, left) and after coordinate transformation, a yellow laser beam was applied to the corresponding area on the fish head (Fig. 6b, right). The full x width at half-maximum (FWHM) of the photostimulation intensity profile is 7.8 *µ*m and the y FWHM is 6.7 *µ*m (Fig. S7d). We could identify the activation of the corresponding region in the green channel before and after the onset of optogenetic stimulation (Fig. 6d and Fig. S7a-b). During stimulation, we aim to consistently target the same area (Fig. S7c) and Fig. 6e shows the displacement (or error) between the actual position of the light spot and the target position during a 50-second experiment. The shaded red regions indicate periods of swift fish motion during which the yellow laser was deflected out of the FoV to avoid targeting the wrong area. Several factors, including a limited speed to update the position of stimulation (Fig. 6a) and tracking error (Fig. 2), make it challenging to correctly stimulate the targeted area when larval zebrafish were swimming. We therefore decided to perform optogenetic manipulation *only* during inter-bout intervals (i.e., when the animal is relatively still, see Methods).

### 2.5 Optogenetic manipulation of nMLF

Finally, we demonstrate how our integrated system, which combines 3D tracking, brain-wide Ca^2+^ imaging and optogenetic stimulation of a defined brain region, can be used to probe the relationship between neural activity and behavior. We performed a transient (1.5 second) unilateral optogenetic stimulation (Fig. 7a) of the tegmentum region including nMLF (30, 31), which quickly induced an ipsilateral turn within 2 s in the vast majority of cases (VideoS3). Fig. 7b shows example bouts from 1 fish and Fig. 7c plots the turning angle distribution from 6 fish. Spontaneous turns in the absence of light stimulation did not show directional bias, and the magnitude of a turn was smaller (Fig. 7b-c). As a control, when the same light stimulation was applied to zebrafish that did not express the light-sensitive protein ChrimsonR, the animals’ movements did not show directional preference: they exhibited more forward runs and when they turned, the turning amplitude was much smaller (Fig. 7d-e).

**Figure 7.**
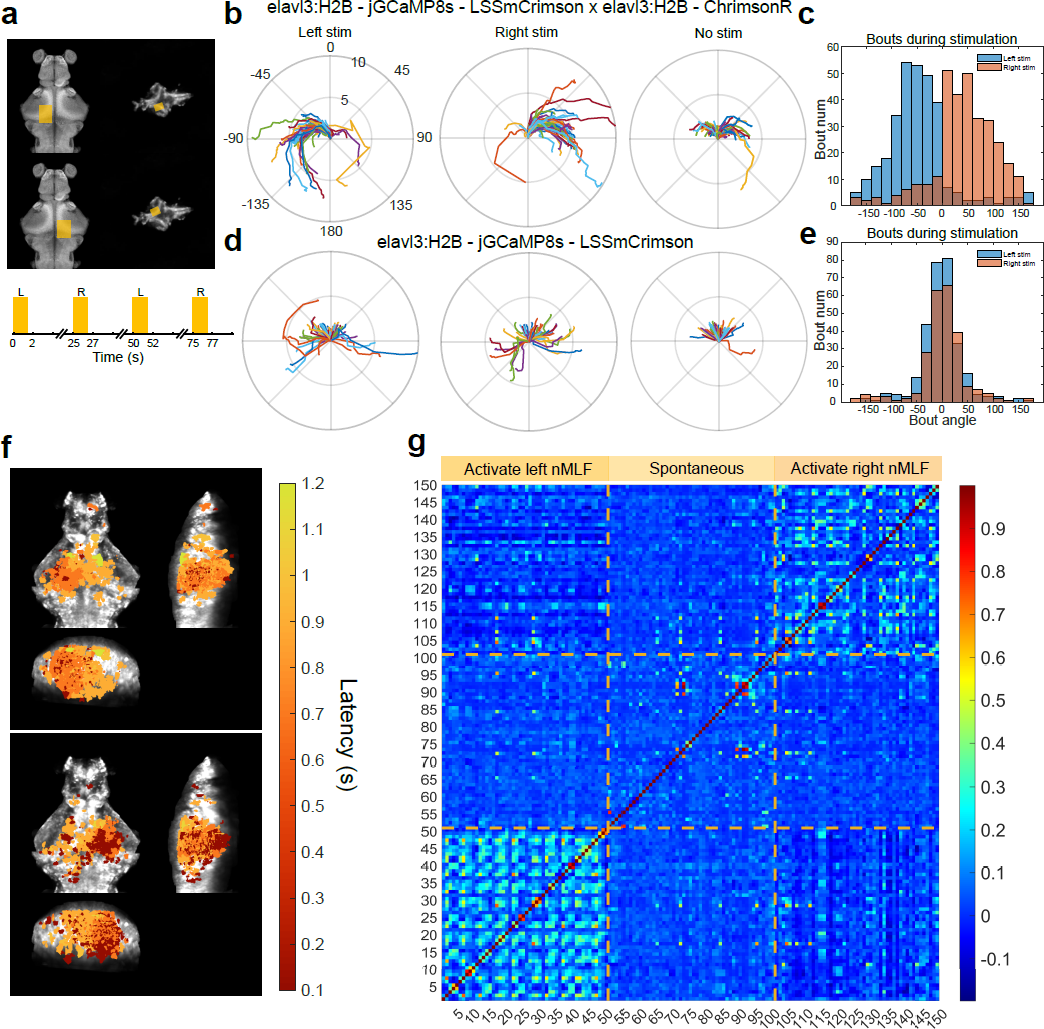
Optogenetic activation of the unilateral nMLF region induced ipsilateral turning behavior and activity changes. **a.** Stimulation regions and paradigm. The left and right mid-brain regions including nMLF were alternatively stimulated (VideoS3). **b.** Example bout trajectories from a elavl3:H2B-jGCaMP8s - elavl3:H2B-LSSmCrimson *×* elavl3:ChrimsonR-tdTomato fish. **c.** Histogram of Bout angle from all elavl3:H2B-jGCaMP8s - elavl3:H2B-LSSmCrimson *×* elavl3:ChrimsonR-tdTomato fish (n = 6). **d.** Example bout trajectory from a elavl3:H2B-jGCaMP8s - elavl3:H2B-LSSmCrimson fish without the expression of opsin (ChrimsonR) in neurons. **e.** Histogram of bout angle from all elavl3:H2B-jGCaMP8s - elavl3:H2B-LSSmCrimson fish (n = 5, Methods). **f.** Activity appeared after optogenetic stimulation of the unilateral nMLF region. Active regions were color coded based on their response onset time when the corrected signal intensity reached 20% of their maximum after the optogenetic manipulation. **g.** Pairwise Pearson’s *r* of brain-wide activity between time frames, defined as 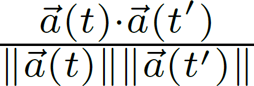. Time frames include 500-millisecond periods (5 frames in each trial and 20 trials) after activating the unilateral nMLF region, as well as randomly selected 500-millisecond epochs when no optogenetic manipulation was performed.

All-optical interrogation enabled us to investigate how local optogenetic manipulation impacts brain-wide activity in a freely swimming zebrafish. Fig. 7f reveals the appearance of neural activity across brain circuits after optogenetic activation of nMLF. The trial-averaged active regions were colored according to their response onset time, when the activity amplitude reached 20% of its maximum after optogenetic manipulation. We found that evoked neural activity appeared in different regions of the brain, including the tegmentum, the optic tectum, the torus semicircularis, and the cerebellum (Fig. S8). In particular, a total of *n* = 1788 ROIs exhibited prominent Ca^2+^ activity during optogenetic stimulation of the left or right nMLF region. The population neural activity pattern, which can be viewed as an n-dimensional vector 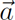(t), exhibited higher Pearson’s correlation coefficient across trials when unilateral stimulation was applied to the same side of the brain than when stimulation was applied to the opposite side (Fig. 7g and legends).

## 3 Discussion

Freely-swimming zebrafish exhibit different internal states and behavioral responses to sensory inputs than head-fixed zebrafish (17, 31). By combining robust tracking in different conditions, Ca^2+^ signal correction based on dual color fluorescence imaging, and optogenetic manipulation of freely swimming zebrafish, our method enables more accurate read-outs of brain activity associated with different behaviors, and all-optical interrogation of the brain-wide circuit in a freely swimming larval zebrafish (Fig. 8).

**Figure 8.**
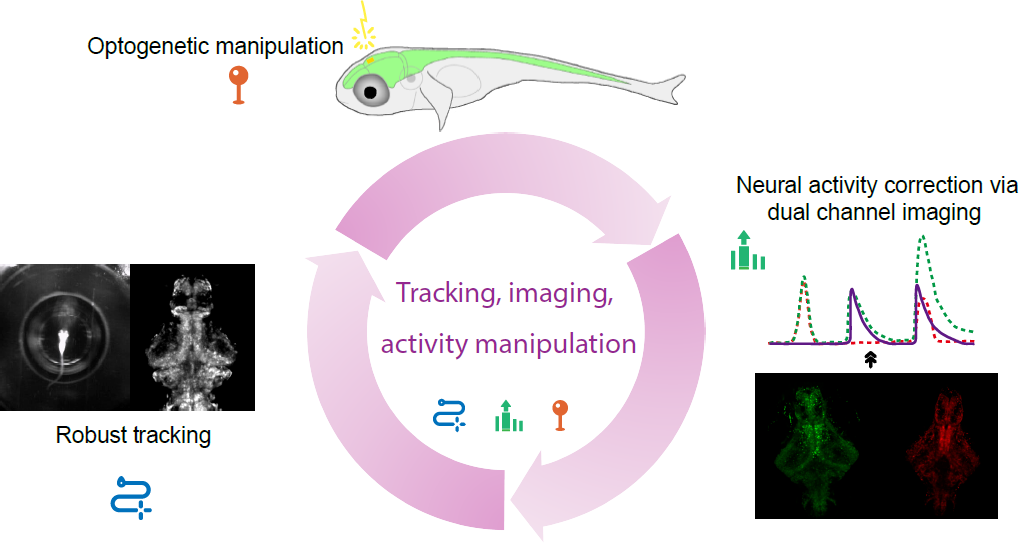
Summary of the all-optical interrogation method. Robust tracking, accurate extraction of Ca^2+^ activity, as well as manipulation of neural activity in specific brain regions enable us to investigate the neural mechanisms underlying different behaviors of freely-swimming zebrafish.

Extracting neural activity from Ca^2+^ signals in the presence of strong noise is a challenging task. Here, we tackled this problem employing the AF method, which has been widely applied in other biomedical signal processing applications (32, 33). Adaptive filters can effectively eliminate motion artifacts and interference from strongly correlated noise. Compared to other motion correction methods (34), the AF algorithm is fast and does not require prior knowledge. These advantages make it more suitable for closed-loop online experiments.

Improving the correlation and signal-to-noise ratio (SNR) of the green and red signals allows our AF algorithm to obtain more accurate signal correction results (Fig. S5). Using less pigmented and more transparent zebrafish, such as the casper line (35) for brain imaging, helps to reduce the chromatic aberration caused by pigmentation and tissue scattering and could thus improve the correlation between red and green signals. Using brighter red fluorescent proteins and Ca^2+^ indicators with a larger dynamic range (36) would further improve the SNR and thus make the extraction of the activity signal more accurate. Another possibility is to use SomaGCaMP (37) instead of nuclear localized GCaMP, which would increase the amount of fluorophore expression.

The AF algorithm is not always accurate for all brain regions, especially when the correlation and SNR of the dual-channel signals are low. To address this issue, we may need to develop refined models that use the statistics of dual channel signals. One possible improvement is to perform an in-depth statistical analysis of the data captured from freely swimming EGFP × LSSmCrimson zebrafish. We could also extract features of neural activity-induced signal changes in jGCaMP8s x LSSmCrimson zebrafish paralyzed by bungarotoxin. These prior information, when combined with the location of each brain region, could be used to develop a more accurate model for dual-channel signals.

Previous zebrafish optogenetic experiments can be divided into two categories: (1) optogenetic manipulation of specific brain regions with spatially patterned illumination in head-fixed zebrafish (12, 14); (2) manipulation of freely swimming zebrafish with spatially localized photosensitive proteins using full-field light (38, 39). The first method can provide spatially accurate stimulation, but behavior responses are restricted and not natural. The second approach allows optogenetic manipulation of naturalistic behavior, but generating fish lines expressing photosensitive proteins in desired brain regions is challenging and only one spatial pattern of stimulation can be applied to the fish. Our system aims to overcome the limitations of both approaches: it allows selective light stimulation of specific brain regions and rapid switching of stimuli between multiple regions in freely swimming zebrafish.

The current optogenetic module uses a laser without beam expansion, and manipulation of a selected brain region is achieved by high-speed 2D galvo mirror scanning, a design that greatly reduces the loss of laser energy and allows for ultra-high intensity of light projection. An alternative method is to generate patterned illumination using the digital micromirror device (DMD) (13, 40), which could be less energy efficient. However, our design results in optogenetic manipulation without Z-resolution (Fig. 6c). A set of lenses could be added before the galvo mirror for beam expansion. This would allow the beam to converge only near the focal plane, and the rapid decrease in light intensity away from the focal plane would enable the activation of neurons only near the focal plane. The position of the focal plane could be adjusted by moving the lens with a piezo.

Our real-time optogenetic system updates its illumination pattern at 10 Hz. The main factor that limits the speed of the system is the time it takes to reconstruct the volumetric image of the fish that is used to register in the ZBB atlas. Using deep learning algorithms (41–44) instead of the traditional Richardson-Lucy iterative reconstruction method is expected to greatly accelerate the speed of reconstruction and make our system more responsive to rapid and high-frequency head movements.

We use visible light for single-photon optogenetic manipulation, an approach that allows us to manipulate a large range of brain regions nearly simultaneously and has minimal thermal effects compared to infrared light. However, visible light is easily scattered by brain tissue, making manipulation less spatially accurate, especially for deep brain regions. Recently developed two-photon optogenetics and holographic technique have enabled manipulation of multiple neurons at different locations in a fixed 3D volume (9, 12). Two-photon microscopy has also demonstrated its ability to track and stimulate a single neuron in a freely moving Drosophila larva (45). The inherent nature of two-photon excitation can significantly reduce tissue scattering and improve manipulation accuracy (46). Combining two-photon optogenetics with one-photon volumetric imaging in freely swimming zebrafish is a promising future direction if the spatial accuracy of optogenetic manipulation at the single-neuron level is critical; if speed and cost are critical, then single-photon optogenetics would be the better choice.

Molecular biology approaches can be used to further improve the accuracy and adaptability of optical read-outs and the manipulation of defined neural populations. For example, the integration of novel nuclear localization sequences optimizes the confinement of calcium indicators and RFP within the cell nucleus (47). This optimization could intensify the sparsity of fluorescence expression, thereby elevating the resolution of the reconstructed images. With the aid of suitable promoters and the GAL4 / UAS system, opsins can be expressed in defined brain regions or cell-types (48–50). This capability would enable us to explore the influence of anatomically and/or genetically defined cell assemblies on brain-wide activity and animal behavior.

In conclusion, we anticipate that our all-optical technique, when combined with recent development in volumetric imaging methods (17, 19, 51–57), would significantly advance the investigation of neural mechanisms underlying various naturalistic behaviors in zebrafish and other model organisms (25, 58–61).

## 4 Methods

### 4.1 Hardware

The new system was an update of XLFM (16). The upgraded system consists of three main components: a 3D tracking module, a dual-color fluorescence imaging module, and an optogenetic manipulation module.

The 3D tracking module used a high-speed camera (0.8 ms exposure time, 340 fps, Basler aca2000-340kmNIR, Germany) to capture the lateral motion of the fish. We developed a U-Net (21) based real-time system that could rapidly identify the head and yolk position. The error signal between the actual head position and the set point was then fed into the tracking model to generate output signals and control the movement of a high-speed customized stage. The autofocus camera (100 fps, Basler aca2000-340kmNIR) behind a 5-microlens array captured 5 images of the fish from different perspectives. The z position of the fish can be estimated by calculating the inter-fish distance based on the principle of LFM. The error signal between the actual axial position of the fish head and the set point was then fed into the PID to generate an output signal to drive a piezo (PI P725KHDS, 400 µ m travel distance) coupled to the fish container.

In the dual-color fluorescence imaging module, a blue excitation laser (Coherent, Sapphire 488 nm, 400 mW) was expanded and collimated into a beam with a diameter of ∼ 25 mm. It was then focused by an achromatic lens (focal length: 125 mm) and reflected by a dichroic mirror (Semrock, Di02-R488-25X36, US) in the back pupil of the imaging objective (Nikon N25X-APO-MP, 25X, NA 1.1, WD 2 mm, Japan) resulting in an illumination area of ∼ 1.44 mm in diameter near the objective focal plane. This 488 nm laser was used to simultaneously excite jGCaMP8s and LSSmCrimson. Along the fluorescence imaging light path, the fluorescence collected by the objective was split into two beams by a dichroic mirror (Semrock, FF556-SDi01-25X36) before the microlens arrays and entered the two sCMOS cameras (Andor Zyla 4.2, UK) separately. Each lenslet array consisted of two groups of microlenses with different focal lengths (26 mm or 24.6 mm) in order to extend the axial field of view while maintaining the same magnification for each subimage. Both lenslet arrays were conjugated to the objective back pupil by a pair of achromatic lenses (focal lengths: F1 = 180 mm and F2 = 160 mm). Two bandpass filters (Semrock FF01-525/45 and Semrock FF02-650/100) were placed before 2 cameras, respectively, to block light from other wavelengths.

A 588 nm laser (CNI MGL-III-588, China) reflected by a 2D galvo mirror system (Thorlabs GVS002, USA) was used for optogenetic manipulation. The midpoint of two galvo mirrors was conjugated onto the back pupil of the imaging objective by a pair of achromatic lenses (focal lengths: F1 = 180 mm and F2 = 180 mm). A dichroic mirror (Semrock, Di01-R405/488/594-25X36) reflected the 588 nm laser and transmitted the green and red fluorescence.

### 4.2 A U-Net neural network to track fish position and orientation

We used a simplified U-Net (21) model to detect the head and yolk of the fish. The model contains only two downsampling layers and two upsampling layers, which improves the detection speed. The heat map is used as the output of the model, showing the location of the target point and the confidence level.

To create a training dataset, we used custom MATLAB tools (Natick, MA). First, we used k-means to extract the key frames in the video. The key frames covered the variety of fish movements and the complexity of the background. Second, we read the first three keyframes of each video and manually marked the position of the fish’s head and yolk. We rotated the three keyframes until the fish’s head was facing the same direction, calculated the average image, and rotated the average images at 5-degree intervals to create 72 templates. Then, the positions of the head and yolk of the fish in all keyframes were determined using template matching. We manually checked the position marked on each keyframe and corrected for the wrong label. Finally, we created a training dataset by combining all labeled frames and dividing them into subsets of test and training frames.

We used pytorch to build, train, and test the model. For training, we used the Adam optimizer and MSE loss function, with a batch size of 32, an initial learning rate of 0.001 and a gradual decay. We saved the model with the lowest loss in the test set and stopped training when the loss was no longer decreasing. We implemented the models in our tracking system using TensorRT. We reduced the model precision to Float16 to improve inference speed without sacrificing inference accuracy.

### 4.3 Model predictive control (MPC)

We adopted the MPC method in (17) to control the X-Y motorized stage. We modeled the motion of the stage and the fish, and then selected the optimal stage input by minimizing future tracking error. The stage was modeled as a linear time-invariant system, whose velocity was predicted by convolving the input with the impulse response function of the system. The motion of the fish during a bout was modeled as a uniform linear motion. Instead of directly predicting the trajectory of the fish brain, as in (17), we first predicted the trajectory of the fish yolk, which is much straighter, especially at the beginning of a bout. We then predicted the fish brain position by shifting along the current heading vector. The loss function to be minimized is the sum of squares of tracking error over six time steps into the future, plus an L2 penalty for stage input. We replaced the L2 penalty on the future planned acceleration vector (17) with the L2 penalty on stage inputs, which empirically reduced stage vibration.

### 4.4 Volumetric image registration

Accurate alignment of volumetric images of freely swimming zebrafish larvae is essential for subsequent signal correction and neural activity analysis. Because the green and the red fluorescence signals are derived from splitting a single beam of light, the raw reconstructed volumetric images between the two channels differ only by a simple affine transformation. The red channel is activity-independent and the intensity difference between frames is relatively smaller than that in the green channel. Therefore, we first performed multistep alignment on the red channel and then applied identical operations on the green channel (Fig. 3a). Direct alignment of raw images is time-consuming and data-intensive. To address these challenges, we designed and implemented the following four-step registration pipeline.

- **Crop**. We first rotated the reconstructed original 3D images (600 × 600 × 250 voxels) based on the orientation of the fish head recorded by the behavior camera. This rotation aligned the rostrocaudal axis of all frames in the same direction. We then cropped and removed the black space surrounding the fish. Finally, the images were resized to 308 × 380 × 210 voxels, which are the dimensions of the ZBB atlas template (27).
- **Approximate registration**. We selected one cropped image and aligned it with the ZBB atlas to generate a unified template for the whole sequence. Next, we use the CMTK toolbox to register each frame with the template using the affine transformation (22, 62). Using the "correlation ratio, CR" as the registration metric, we achieved the best result (63). With "OpenMP" multithread optimization, we were able to finish one frame of approximate registration in 25 seconds (64).
- **Remove fish eyes**. Because the rotation of fish eyes could seriously affect the next step nonrigid registration, we used a U-net neural network to automatically remove fish eyes.
- **Diffeomorphic registration**. To handle the minute changes in an image caused by breathing, heartbeat, and body twisting, we found it necessary to refine our approximate registration. We used the optical flow algorithm ’Maxwell’s demons’ to implement non-rigid registration in the final phase (23). To increase the precision of the optical flow method with our data, we generated a fresh alignment template. This was achieved by taking an average from every tenth frame across every hundred frames within the image sequence. Subsequently, we lined up each segment of the sequence with the newly created alignment templates. This algorithm was developed using MATLAB and can be speeded up via a GPU. In practice, the execution time for each frame was averaged at 72 seconds on a single RTX 3090 GPU. Efficiency can be further optimized by deploying multiple GPUs.

To evaluate the alignment accuracy, we first implemented our multistep registration pipeline on the red and green channels of a swimming larval zebrafish. The green channel in this case contained the EGFP signals that were sparsely expressed in the neuronal nuclei. We used the MATLAB toolkit "CellSegm" (65) to perform cell segmentation and obtain the centroid coordinates of neurons expressed by EGFP in every frame. We then matched each neuron’s positions in a post-registered time frame to its correspondent in a fixed template frame using the Hungarian algorithm. We identified 164 ±15 (mean ±SD) neurons in all time frames and 195 neurons in the template. Due to uneven distribution of excitation light in the FoV, scattering, and attenuation of fluorescence by brain tissue and body pigments, not all neurons’ cell bodies were visible in every frame. We introduced the searching radius, a hyperparameter in the Hungarian algorithm that controls how far the algorithm will look for matches. The root mean square displacement (RMSD) between the positions of pairs of matched neurons and the matching ratio both increased with the searching radius and plateaued at a large radius (Fig. S2c). We selected the result with a 5-voxel search radius as our best estimate of alignment accuracy (Fig. 3d and Fig. S2c), because it was large enough to find most of the true matches.

### 4.5 Segmentation

After applying multistep registration (section 4.4) to the activity-dependent green channel, we performed cell segmentation based on temporal correlation in the green channel, following the approach introduced in (66). We calculated the average correlation between each voxel and its 14 neighbors to obtain a "correlation map". We then implemented the watershed algorithm on this correlation map to obtain a preliminary segmentation result. For each voxel, we further analyzed its correlation with the average activity of that specific segmented region and obtained the ’coherence map’. Finally, we used a threshold filter on the coherence map to obtain the final segmentation results, which was also applied to the red channel.

The correlation-based segmentation was not suitable for the elavl3:H2B - EGFP × elavl3:H2B - LSSmCrimson fish data because the green channel lacks the neural activity signals. Instead, we divided the entire image into 8 × 8 × 6 voxel regions. The grid size was close to the mean segmented ROI size of our Ca^2+^ imaging data.

### 4.6 Normalized least-mean-square (NLMS) adaptive filter

We start by presenting a phenomenological model of the fluctuation of fluorescence signals in the activity-dependent green channel and the activity-independent red channel. Noise in the red channel has two major contributions: motion-induced fluctuation of fish and independent noise introduced along the optical pathway and by the sCMOS camera. Here we model the red LSSmCrimson signal *r*(*n*) from a given ROI at time frame *n* as

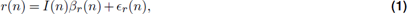

where *ɛ*_*r*_(*n*) is independent noise, *I*(*n*) is the local excitation light intensity and *β*_*r*_(*n*) is the baseline fluorescence, which in theory only depends on the number of fluorophores expressed in the neuron. However, fish movements in 3 dimensions make both *I*(*n*) and *β*_*r*_(*n*) time dependent.

We model the green jGCaMP8s signal *g*(*n*) from the same ROI in a similar way by incorporating the Ca^2+^ activity *a*(*n*)

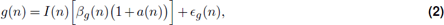

where *ɛ*_*g*_(*n*) is independent noise and *β*_*g*_(*n*) is the baseline. If the emission light fields of the green and red fluorophores captured by the imaging objective and the camera are identical, then we can identify *β*_*g*_(*n*) = *αβ*_*r*_(*n*), where α is a proportionality constant. Therefore, a simple division between the green and red signals is sufficient to extract Ca^2+^ activity, provided that the independent noise is small. In practice, due to various scattering of brain tissue and obstruction of body pigments, we are agnostic about the relationship between *β*_*g*_(*n*) and *β*_*r*_(*n*); but it is reasonable to assume that they are strongly correlated in a complicated and time-dependent way, which is consistent with our observation and analysis of the raw EGFP and LSSmCrimson signals.

Under the assumption that motion-induced fluctuations are strongly correlated between the green and the red channel, and such fluctuation is history-dependent, we decided to use an adaptive filter (AF) to extract Ca^2+^ activity corrupted by motion artifacts. We aim to predict the green channel signal *g*(*n*) from the red channel *r*(*n*) using the normalized least mean squares (NLMS) AF algorithm (67). The NLMS algorithm continuously subtracts 4.7 Galvo voltage matrix the predicted signal *ĝ*(*n*) from *g*(*n*) so that the residue *e*(*n*) = *g*(*n*) − *ĝ*(*n*) is minimized. The predicted *ĝ*(*n*) is a weighted sum of the history-dependent red signal:

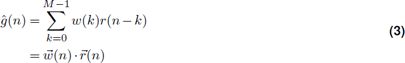

where *M* is the filter length and here we set *M* = 2. We iteratively update the weight (i.e., filter) vector 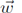 using gradient descent:

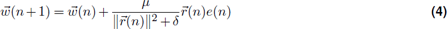

We follow (68) to determine the iteration step size *µ*. Once *ĝ*(*n*) was computed, we made the following assumption that:

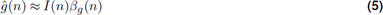

As a result, the Ca^2+^ activity is given by

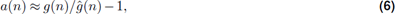

provided that ɛ_*g*_(*n*) ≪ *ĝ*(*n*), namely the independent noise is much smaller than motion-induced fluctuation.

#### Performance of AF algorithm

To identify the factors that affect the inference ability of the AF algorithm, we added randomly timed synthetic signals with fixed amplitude to the EGFP signals of freely swimming zebrafish. We then evaluated the AF inference performance by calculating the cross-correlation (Pearson’s *r*) between the AF inferred signal and the synthetic signal for each ROI. We visually represented each ROI’s correlation as colored scattered points in two dimensions with the correlation between dual-channel signals on the y-axis and the coefficient of variation (CV) of the red channel on the x-axis for different synthetic signal amplitudes (Fig. S5, left column). ROIs with a higher correlation between dual-channel signals and smaller CV typically exhibit better AF inference performance. As the amplitude of the synthetic signal increases (defined as the CV of an ROI’s EGFP signal multiplied by a constant), the overall inference performance also increases (Fig. S5, right column).

To establish a criterion for screening ROIs with a potential high AF inference performance, we binarized the *r* between AF inferred signal and synthetic signal with a threshold value of 0.5. We considered ROIs with a *r* greater than 0.5 to have a good inference performance (Fig. S4). We then used polynomial logistic regression to perform a binary classification task, which can be modeled as:

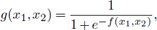

where *f* is a cubic polynomial function with fitted coefficients, and *x*_1_, *x*_2_ are CV of the red channel and Pearson’s *r* of dual-channel signals respectively. For visualization purposes, the pink regime in Fig. S5 corresponds to *g* > 0.7 while the blue one corresponds to *g* < 0.7; the dashed line dividing the two regimes can be viewed as a decision boundary. The area under the ROC curve (AUC) was used to quantify the classification performance.

We found that the measured medium amplitude of jGCaMP8s in head-fixed larval zebrafish was about 4 times the CV of EGFP signals in freely swimming zebrafish (Fig. S3). We therefore selected the fitted model when the amplitude of synthetic neural activity is four times (4×) the CV of the EGFP signals (Fig. S5c). Note that when the same model (Fig. S5c) was applied to synthetic signals whose amplitudes were sampled from a defined statistical distribution instead of a fixed amplitude, we found a similar classification performance.

### 4.7 Galvo voltage matrix

We placed a fluorescence plate under the microscope objective and adjusted the stage height and laser intensity to maximize the brightness and minimize the size of the fluorescence spot excited by the optogenetic laser. The galvo input is a voltage pair (GalvoX, GalvoY). We varied the galvo voltage input with an interval of 0.1 V and a range of −1.5 V to +1.5 V in both the X and Y directions. We recorded 10 frames per voltage pair. The raw recorded images were resized to 512 × 512 pixels and reconstructed. We took the coordinates of the brightest point on the image of each voltage pair. If there was no bright spot in the FoV, the voltage pair was deleted. As a result, the correspondence between some of the galvo input voltage pairs and image coordinates was known. Assuming a linear transformation relationship between the voltage pairs and the coordinates, we found the affine transformation matrix using the known points. Then, we calculated the galvo voltage pair corresponding to each point in the image and stored it as the GalvoX and GalvoY voltage matrices.

### 4.8 Online optogenetics pipeline

A custom C++ program was used to implement the optogenetic system. After the images were captured by the red fluorescence camera, the image reconstruction and alignment process were implemented in the GPU, while the coordinate transformation and the control of the galvo were implemented in the CPU.

The image processing algorithm was run on an RTX 3080 Ti GPU using CUDA 11. We resized an image from 2048 × 2048 pixels to 512 × 512 pixels using the AVIR image resizing algorithm designed by Aleksey Vaneev (https://github.com/avaneev/avir). Due to the reduced image size and memory consumption, we could use the PSF of the whole volume to do the deconvolution with a total of 10 iterations. The size of the reconstructed 3D image is 200 × 200 × 50 voxels. It took about 75 ms to reconstruct one frame.

We used TCP to communicate between the tracking system and the optogenetic system. We rotated the fish head orientation of the 3D image to match that of the ZBB atlas using the fish heading angle provided by the tracking system. We then found the maximum connected region by threshold segmentation and removed the redundant pixels outside the region. The size of the image after cropping was 95 × 76 × 50 pixels, which is the same as the ZBB atlas. Finally, we aligned the 3D image with the standard brain by affine transformation using a transformer neural network model. The rotation, cropping, and affine alignment took about 10 ms.

The coordinate transformation first calculated the inverse of the affine matrix and the rotation matrix. The user-provided coordinates of the region on the ZBB atlas were then multiplied by the transformation matrix. Finally, the transformed coordinates were shifted by the upper left corner coordinates of the cropped image. This converted the coordinates of the specified region selected in the ZBB atlas to the coordinates of the actual fish brain.

The voltage pairs to be applied to Galvo were read from the GalvoX and GalvoY voltage matrices (see section 4.7). The voltage signals were then delivered to the 2D galvo system (Thorlabs GVS002, US) using an I/O Device (National Instruments PCIe-6321, US). The galvo system converted the voltage signals into angular displacements of two mirrors, allowing rapid scanning of a specified area.

To avoid targeting the wrong brain region, we decided to deliver light stimulation only during the inter-bout interval. We maintained a queue of length 50 that stored the fish heading angle from the tracking system. The average of the heading angle of the fish in the queue was calculated. If the difference between the received fish heading angle and the average in the queue was greater than 5 degrees, the fish was considered to have entered a bout during the delay, and the laser beam was deflected out of the field of view.

### 4.9 Spatial accuracy of optogenetic stimulation

We selected an ROI of 1 × 1 pixel to test spatial precision during continuous optogenetic stimulation in freely moving zebrafish. A notch filter (Thorlabs NF594-23) was used to exclude the scattering light from the 588 nm laser in Fig. 6c-d. This filter was removed in order to accurately observe the position of the laser in Fig. 6e. It is important to point out that our current optogenetic module does not possess a Z-resolution (see Discussion for potential improvement).

We computed pairwise voxel intensity difference (Δ*F*/*F*) between the average green channel image before stimulation and that during stimulation. Voxels with a value less than 20 were considered noisy background and set to 0. We show a single X-Y and Y-Z plane in Fig. 6d.

### 4.10 Bout detection and behavior analysis

We use the displacement of the tracking stage *δs*(*t*) between consecutive frames to detect the movements of the zebrafish, and create a binary motion sequence by thresholding *δs*(*t*) at 20 µm. To identify the bouts, we merged adjacent binary segments to obtain a complete bout sequence. To characterize bouts, we calculated the angle between the heading direction at the beginning of a bout and the heading direction of each frame during a bout (Fig. 7b and Fig. 7d). The bout angle (Fig. 7c and Fig. 7e) was defined as the angle between the heading direction at the start of a bout and at the end of the bout. Positive values represent right turns.

### 4.11 Brain activity analysis during optogenetic stimulation

Whole brain neural activity were obtained from inferred signals using the AF algorithm, and unreliable regions of the brain were excluded from further analysis (section 4.6).

To characterize brain-wide activity evoked by optogenetic stimulation, we first identified brain regions with significantly elevated neural activity immediately after optogenetic activation of the ipsilateral nMLF. We compared Ca^2+^ activity in these regions with activity at times without optogenetic manipulation and used a rank sum test with multiple comparison correction to identify significant differences. We then calculated the mean activity trace for each ROI by averaging over multiple trials. Finally, we calculated the time (latency) it took for each ROI to reach 20% of its maximum activity. The latency for each ROI is colored in Fig. 7f.

To characterize brain-wide activity and their differences during optogenetic-induced turns, we used a rank sum test with multiple comparison correction to compare Ca^2+^ activity from 3 frames before to 3 frames after the start of a bout and Ca^2+^ activity at other moments. We identified 1788 ROIs that exhibited significantly elevated calcium activity during turns.

We next selected 5 frames of data after the onset of unilateral nMLF photostimulation: 10 trials during left stimulation and 10 trials during right stimulation, for a total of 100 frames. We also selected 10 time segments, each of which contains 5 frames (50 frames in total), randomly selected from the remaining frames without optogenetic manipulation. We calculated the *Pearson’s r* between every pair of the population activity vectors. The results were represented by the similarity matrix (Fig. 7g) using the frame index.

## Code and data availability

The source code can be publicly accessed at https://github.com/Wenlab/OptoSwim.

## Supplementary information

**S1 Supplementary figures.** S1 comprises 8 supplement figures.

**S2 Supplementary movies.** S2 comprises 3 supplement movies.

## ACKNOWLEDGEMENTS

QW was supported by the National Science Foundation of China (NSFC) under grant NSFC-32071008 and STI2030-Major Projects 2022ZD0211900. The authors thank Dr. Kai Wang, Lin Cong, Zhenkun Zhang, and Zeguan Wang for their assistance with the imaging hardware design. The authors thank Dr. Jiulin Du for kindly sharing the elavl3:ChrimsonR-tdTomato fish line. We thank Dr. Jie He and Xiaoying Qiu for their assistance in generating and rearing the transgenic fish lines. Parallel CPU computing was supported by the Hefei Advanced Computing Center.

---

**Figure S1.**
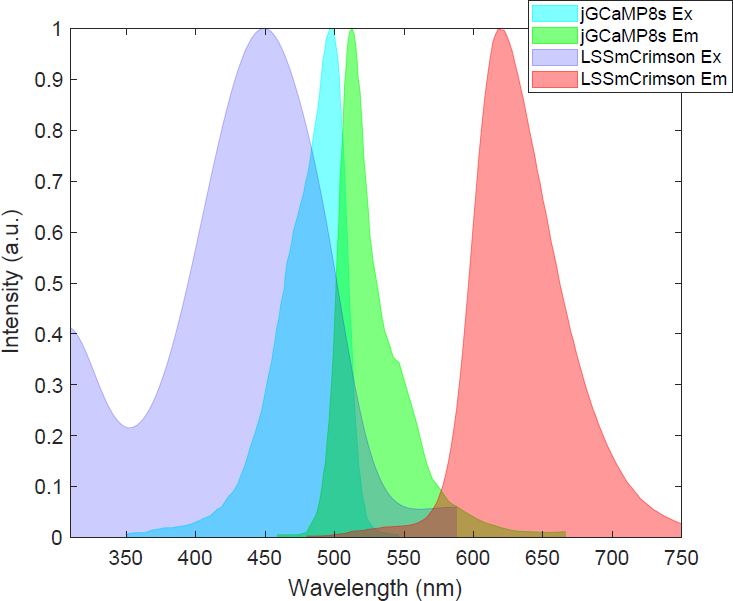
The excitation and emission spectrum of jGCaMP8s and LSSmCrimoson.

**Figure S2.**
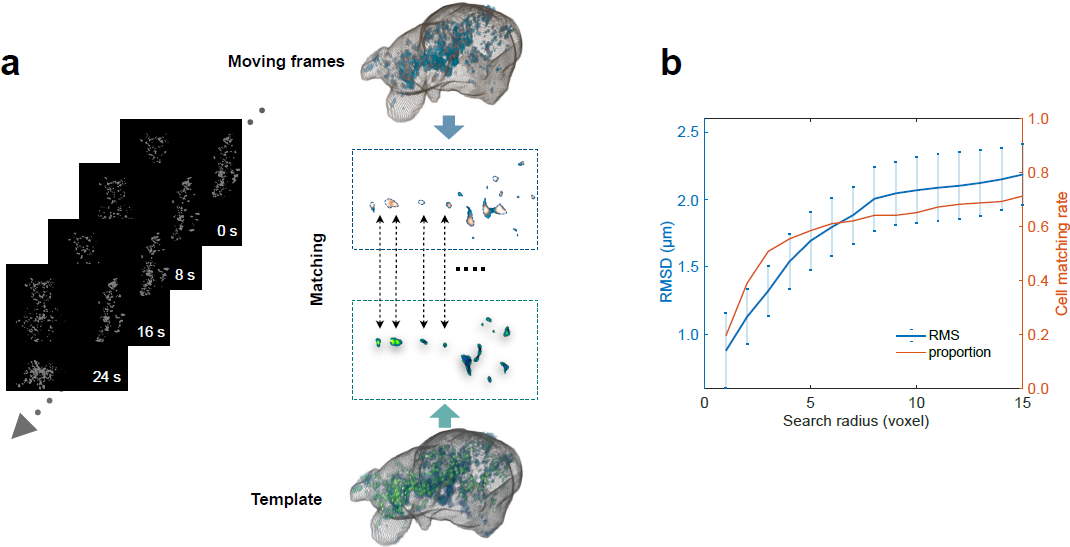
Characterization of the accuracy of registration algorithm. **a.** Perform threshold segmentation on each frame to obtain the centroid coordinates of neurons. **b.** Use the Hungarian algorithm to find the matching neurons between each moving frame and the template. **c.** Compute root-mean-square displacement (RMSD) and neuron matching rate as a function of search radius.

**Figure S3.**
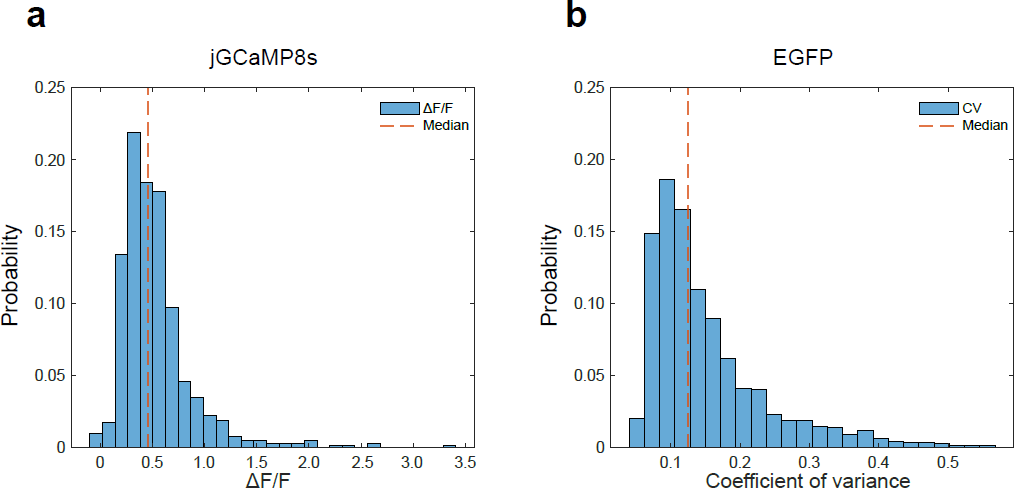
Amplitudes of experimentally measured jGCaMP8s and EGFP signals. **a.** Histogram of Δ*F/F* (baseline to peak) distribution of jGCaMP8s signals in head-immobilized zebrafish. Median = 0.46. **b.** Histogram of coefficient of variation (CV) distribution of EGFP signals in freely swimming zebrafish. Median = 0.13.

**Figure S4.**
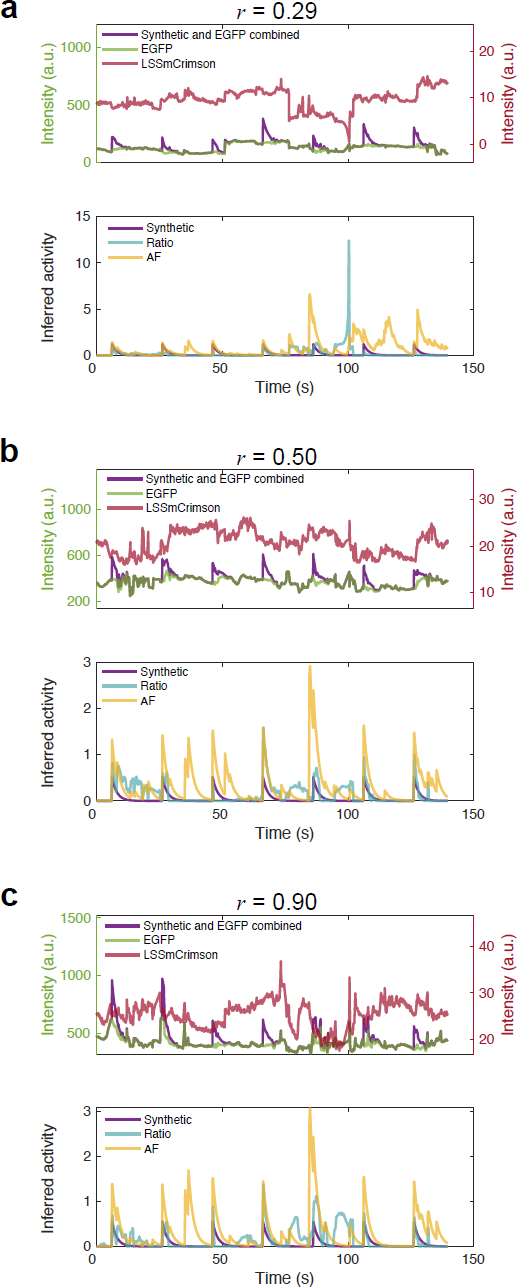
AF inference ability depends on dual-channel correlations and coefficient of variation (CV) of red channel signals. We use the correlation coefficient between AF inferred signals and synthetic signals to measure the inference performance. The following are typical situations from three ROIs: **a.** *r* (Inferred, Synthetic) < 0.5, *r* (Red,Green) = 0.30, and CV(Red) = 0.23. **b.** *r* (Inferred, Synthetic) = 0.5, *r* (Red,Green) = 0.54, and CV(Red) = 0.12. **c.** *r* (Inferred, Synthetic) > 0.5, *r* (Red,Green) = 0.13, and CV(Red) = 0.11.

**Figure S5.**
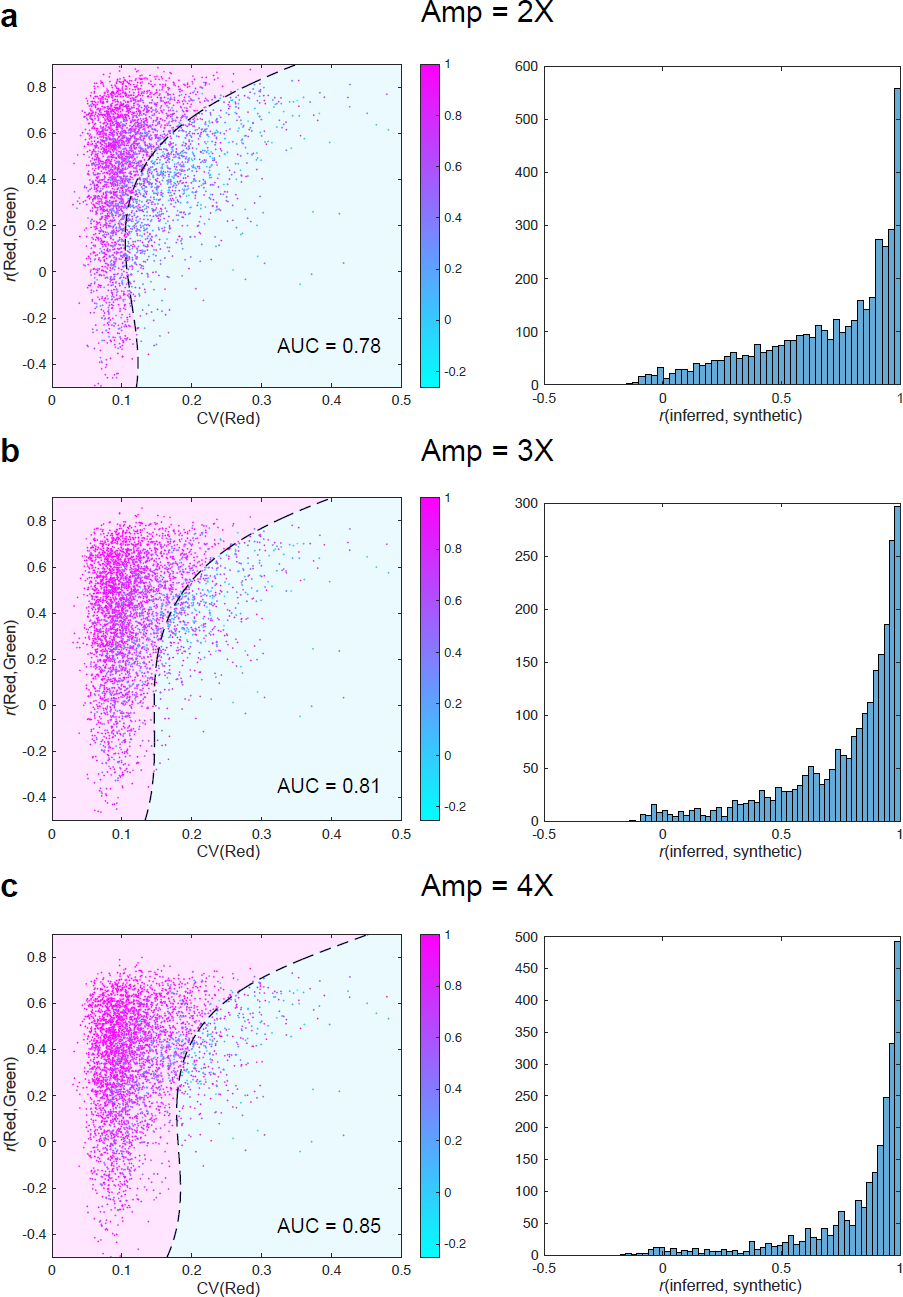
Decision boundary of AF inference performance changes with Ca^2+^ activity amplitude. The amplitudes of synthetic signals are a multiple of the CV of the ROI’s EGFP signal (2*×* in **a**, 3*×* in **b**, and 4*×* in **c**). Left, each panel shows the decision boundary (dashed line), where ROIs (and inferred signals) in the pink regime were accepted whereas those in the blue regime were discarded (Methods). Right, histogram of correlation coefficient *r* between inferred signals and synthetic signals.

**Figure S6.**
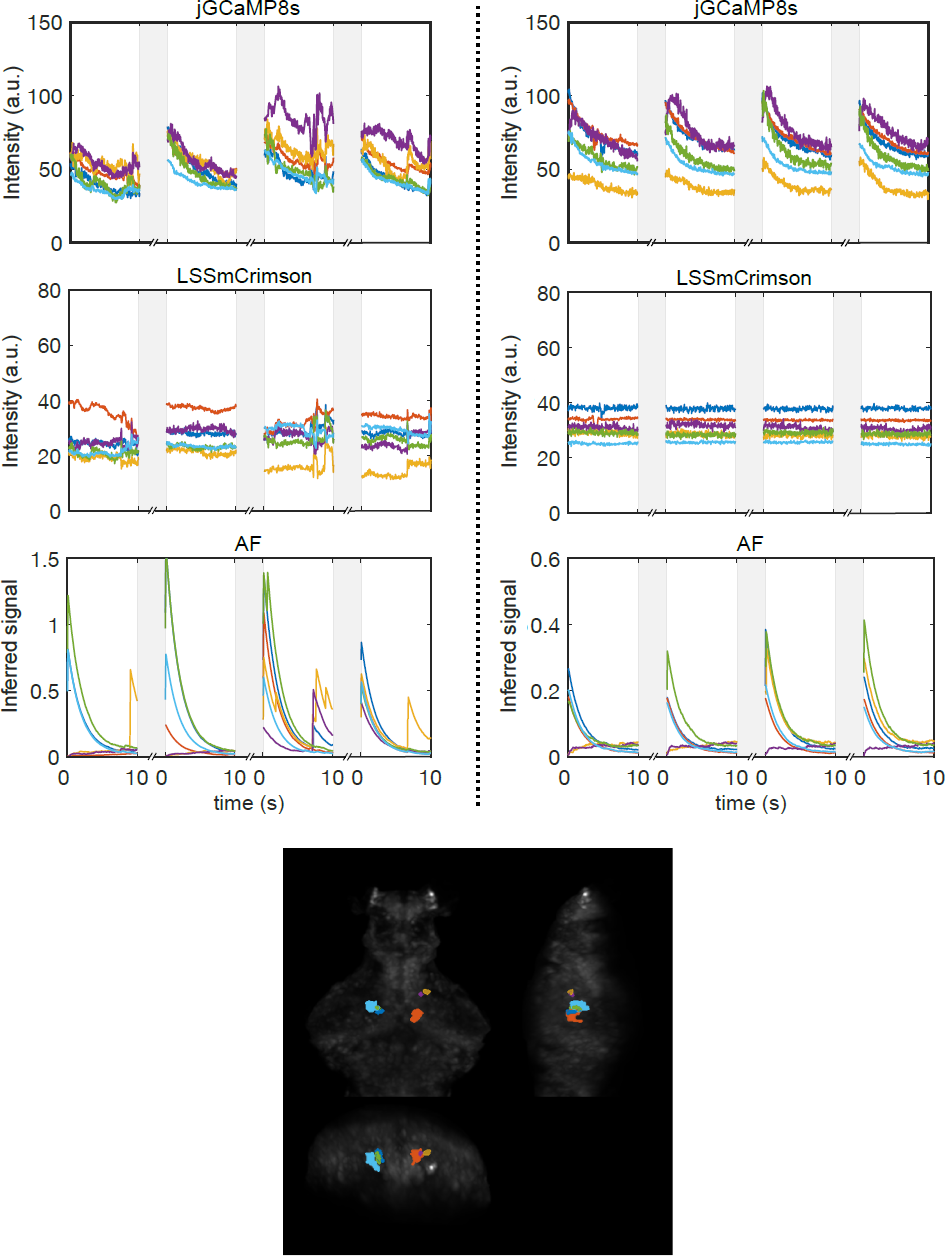
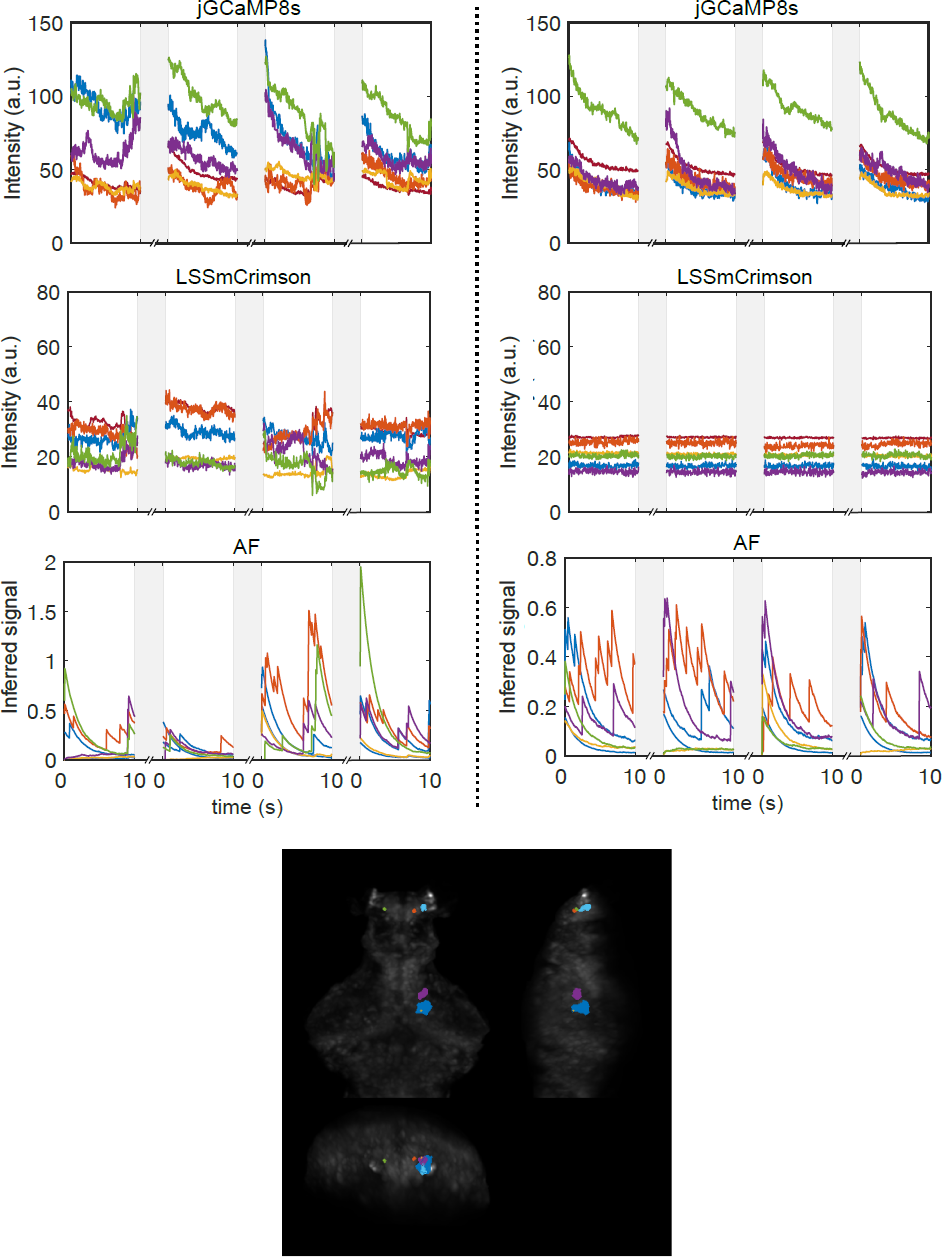
Neural activity in 18 selected ROIs during blue light stimulation (continued on the next page) First and second rows: jGCaMP8s and LSSmCrimson raw fluorescence signals in 6 representative ROIs in which neurons showed prominent Ca^2+^ activity after light stimulus onset. Shaded regions indicate the dark period. Third row: inferred Ca^2+^ activity using the adaptive filter algorithm. Left panels are recordings from freely swimming condition while right panels are from immobilized condition. Bottom: The spatial location of the 6 represented brain regions.

**Figure S7.**
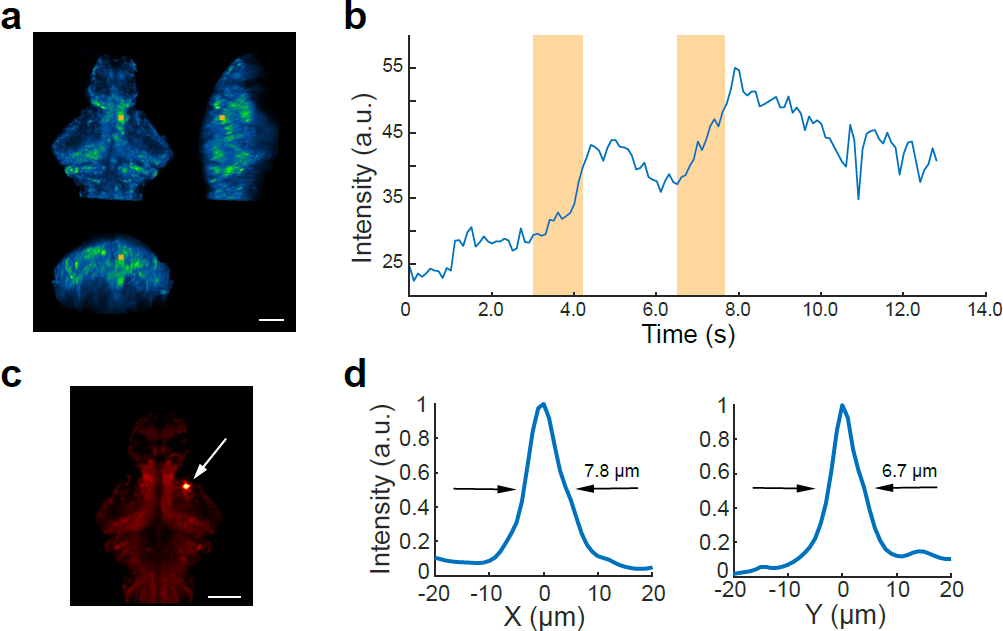
Enhanced Ca^2+^ activity in photostimulated brain region and spatial profile of the laser beam. **a.** Yellow rectangle indicates the stimulated brain region. **b.** The change in fluorescence intensity of the jGCaMP8s in the brain region shown in **a**. Shaded yellow region marked the period of optogenetic stimulation. **c.** Average post-registered fish image (related to Fig. 6e). The white arrow indicates the actual stimulated region during a 50-second experiment in a freely-swimming larval zebrafish. Scale bar, 100 µm. **d.** The measured laser intensity distribution around the stimulated region in **c**. The measured full width at half maximum (FWHM) was 7.8 *µ*m (X axis) and 6.8 *µ*m (Y axis).

**Figure S8.**
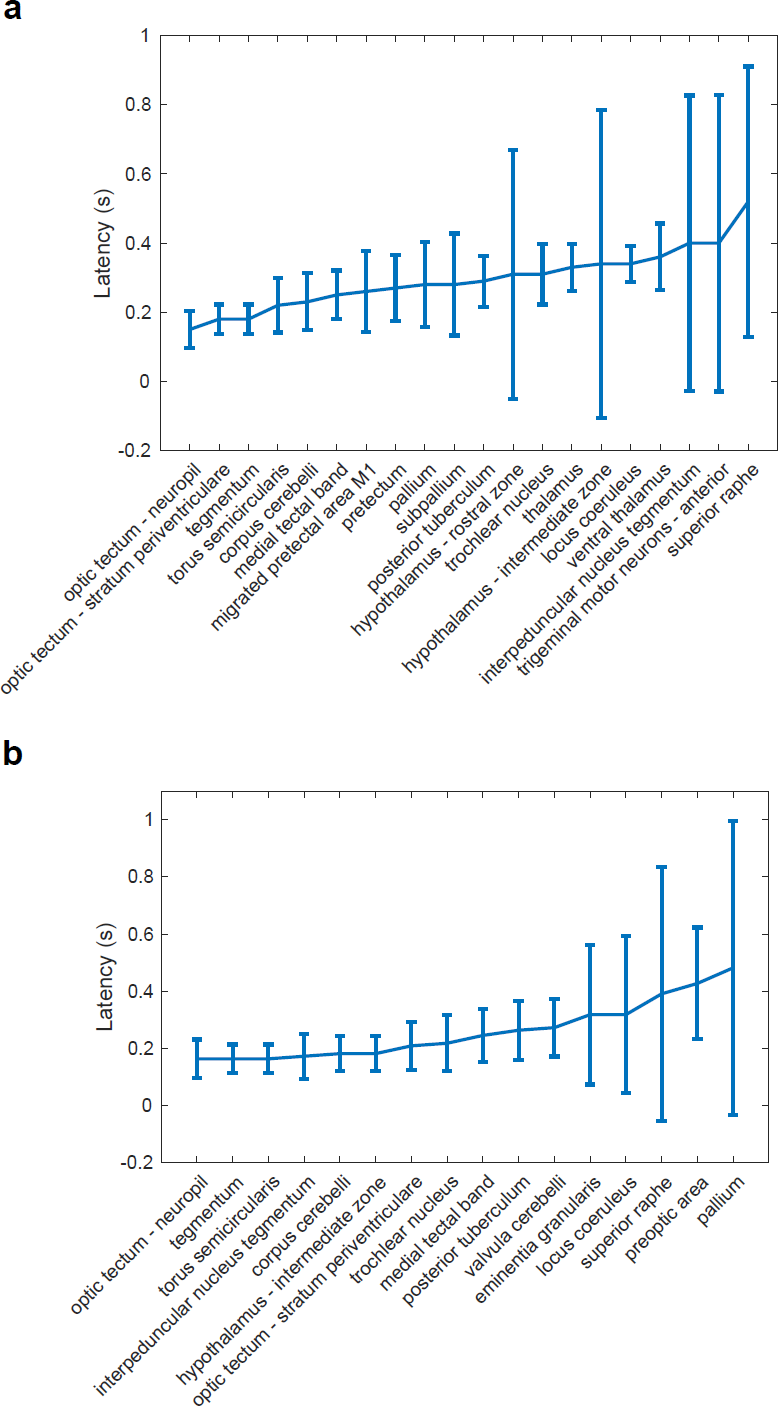
Latency of activity in different brain regions after optogenetic stimulation of unilateral nMLF. The mean onset time of neural activity in each ZBB brain region after optogenetic activation of the left (**a**) or right (**b**) nMLF. The latency was defined as the time at which the amplitude of activity reached 20% of the maximum value of activity. Error bars represent SD.

## Supplementary Videos

### Tracking a freely swimming larval zebrafish

Example NIR tracking video, related to Fig. 2e. A zebrafish was swimming in the presence of water flow. Our tracking algorithm was able to accurately identify the fish head and complete the tracking despite interference from the distracting background.

### 3D registration of a sparsely EGFP-labeled fish brain

Example alignment video, related to Fig. 3b-d. Sparsely EGFP labeled zebrafish were used to test the effectiveness of our alignment method. Left, the fish brain MIP after 3D reconstruction; right, the post-registered fish brain MIP guided by the LSSmCrimson channel.

### All-optical interrogation in a freely swimming larval zebrafish

Example video, related to Fig. 7. On the left, after unilateral optogenetic activation of the tegmentum region containing nMLF, the zebrafish turned ipsilaterally. On the right, changes in whole-brain activity were recorded simultaneously.

